# A high-throughput machine vision-based univariate scale for pain and analgesia in mice

**DOI:** 10.1101/2022.12.29.522204

**Authors:** Gautam S. Sabnis, Leinani E. Hession, Kyungin Kim, Jacob A. Beierle, Vivek Kumar

**Author notes:** Equal Contribution (Gautam S. Sabnis, Leinani E. Hession).

## Abstract

Treatment of acute and chronic pain represent a widespread clinical challenge with poor therapeutic options. While rodents are an invaluable model to study pain, scoring nociceptive responses in clinically relevant paradigms and at high-throughput remains an unmet challenge. Therefore, there is a need for automated, high-throughput methods that sensitively and accurately assess pain and analgesia. Such objective and scalable technologies will enable the discovery of novel analgesics and yield mechanistic insights into the neural and genetic mechanisms of pain. Here, we adopt the open field arena to build a univariate scale for the formalin injection model of inflammatory pain by using a machine learning approach that incorporates 82 behavioral features. This tool outperforms traditional measures of licking and shaking in detection of formalin dose, and was validated using 4 diverse mouse strains. We also detected previously unreported differences in formalin induced nocifensive behaviors that were strain and sex specific. This model also reliably identifies morphine induced antinociception. This novel, sensitive, and inexpensive tool provides a method for quantifying voluntary nociceptive responses to facilitate genetic mapping and analgesic compound screening in a high throughput manner.

## 2 Introduction

In 2013-2014, 40.1% of adult Americans reported suffering from a painful non-cancer health condition, and millions of Americans suffer from chronic pain [1, 2]. In addition to the direct discomfort caused, pain disrupts patient’s lives through comorbidities such as sleep disturbances, mood disorders, and anhedonia [3–5]. The current choice of prescription analgesics are opioids which are accompanied by significant clinical complications like opioid induced hyperalgesia, dose escalation, opioid use disorder, and overdose [6–10]. Pain perception and susceptibility to chronic pain conditions have been found to be heritable, with a number of complex genetic and epigenetic factors involved and better understanding the genetics of pain could lead to novel pain therapies [11–13]. Sex differences in pain thresholds, types of pain, and even analgesic response have been reported in the clinical literature, necessitating appropriate models for pain in both sexes [14–17].

Animal models, particularly mice, are necessary to study the genetic and neurobiological mechanisms of pain, and to test novel treatments. In pre-clinical models, pain can only be inferred from behavior since direct communication of psychological states is not possible. Thus, researchers study the reaction to noxious stimuli, nociception. Nociception is generally measured through operant assays, reflexive evoked measures, or voluntary spontaneous behaviors, each with strengths and weaknesses. Operant assays measure the model organism’s completion of a learned task in response to noxious stimuli or analgesic [18]. As these assays rely on memory and learning, they may not reliably indicate lack of nociception. Reflexive evoked assays typically measure withdrawal reflex [19]. While reflexive evoked measures of nociception are relatively straightforward to collect they have been criticized for having limited relevance to clinical pain conditions [19–22]. Further, the overall failure of reflexive evoked measures to produce successful novel analgesics suggests the need to re-evaluate the dependency of the field on such assays [19–22]. Voluntary non-evoked assays measure freely performed behavior in response to noxious stimuli and represent more naturalistic assays of nociception. Historically, these assays have limited use because they are laborious to manually score and are subject to interobserver reliability. Recent advances in automated behavior extraction have enabled the measurement of non-evoked behaviors at high temporal and spatial resolution in complex environments and include grimace, gait, posture, and home-cage monitoring [19, 21, 23–26]. Here, we seek to use advanced behavioral quantification of nociception to build an univariate pain scale for the laboratory mouse by quantifying spontaneous behaviors.

The formalin test is a well-established assay for localized inflammatory pain wherein the hind paw pad of a mouse is injected with formalin and voluntary behaviors are observed [27]. Commonly, licking/biting of the hind paw is the primary index of nociception. In studies where sex differences in formalin response have been observed, female rodents have been found to show greater sensitivity than males [28]. Composite scores including additional nocifensive behaviors like shaking, favoring, and locomotor activity have been found to be more robust for distinguishing formalin dose [29], suggesting that sampling several modalities of formalin induced voluntary behavior improves sensitivity. These behaviors are labor intensive and difficult to score in mice and are subject to interobserver variability [30–32]. Time sampling while scoring has been used to reduce labor, but are still not high-throughput enough to advance pain and analgesic research [30]. To address these limitations, we previously automated the measurement of licking behavior with a bottom-up camera [32, 33]. However, this specialized arena is small and limits the amount of features, particularly gait, which can be extracted, limiting its sensitivity to detect nociception. Further, defecation and urination obscure the camera view and long-term monitoring with bedding is not possible.

We have developed an open field approach using a top down view for advanced behavior quantification with methods for image segmentation, tracking in complex environments, action detection, pose-based gait, and whole body coordination measurements [34–36]. We have also developed JABS (JAX Animal Behavior System), an active machine-learning interface for training behavior classifiers using pose information [37]. These methods allow us to efficiently and reliably quantify multiple nocifensive behaviors evoked during phenotyping. As scoring multiple behaviors results in more robust nociceptive measures [29], we further expand the measurements taken in the formalin assay to include a richer array of voluntary spontaneous behaviors, many of which have shown relevance in other assays [25]. This system will allow the comparison of specific types of behavioral nocifensive responses, as well as using univariate modeling to more directly compare nociceptive sensitivity across strain and sex, ultimately facilitating studies interrogating the genetic basis of pain sensitivity by increasing the throughput, complexity, and sensitivity of the assay. Additionally, these univariate nociceptive scales can be used to test novel analgesics and to detect sex and strain differences in drug response, improving the throughput and translatability of pre-clinical pain research. Finally, these methods open the possibility for long-term monitoring of nocifensive behaviors in a more naturalistic manner, improving the assessment of chronic pain models, a critically unmet clinical need. To this end, we use males and females from 4 inbred strains of mice previously shown to range from low to high licking response to formalin to asses the sensitivity and validity of our new methods [38]. We also used our approach to assess the behavioral impact of the opioid analgesic morphine on the formalin assay to determine its ability to asses the efficacy of analgesic therapeutics.

## 3 Results

### 3.1 Data collection

In this study we used male and female mice from 4 strains (C57BL/6J, C3H/HeJ, BALB/cJ, A/J) tested with 4 doses of formalin (0.00%/Saline, 1.25%, 2.50%, and 5.00%) in a total of 194 mice. The open field video was processed by a deep neural network based pose estimation network and a tracking network to produce a 12-point pose skeleton and an ellipse fit track of the mouse for each frame (Figure 1A)36, 39]. These per-frame measures were used to train behavior classifiers and engineer features such as traditional open field measures of anxiety, hyperactivity, neural networkbased grooming, and novel gait measures [35, 36, 39]. All features are defined in Table **S1**. The features were used for modeling to produce a univariate pain scale.

**Figure 1:**
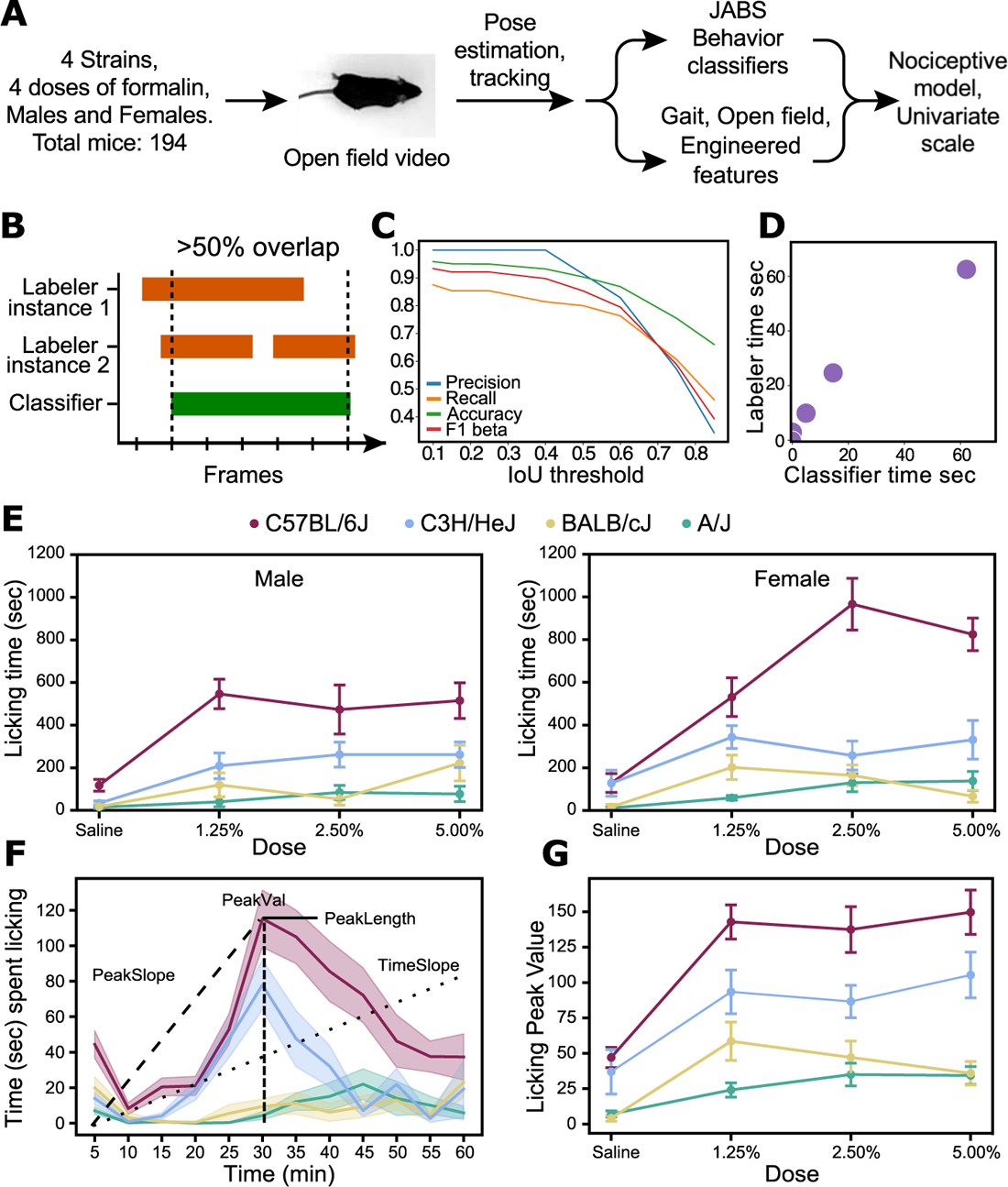
Pipeline and licking classifier validation. A) Pipeline for univariate nociception scale. Mice injected with formalin or saline were recorded in an open field for one hour. Twelve point pose estimation and tracking were extracted. These measures were used to train behavior classifiers and engineer nocifensive features. Features were used to train a model using formalin dose as covariates. B) Illustration of bout-based accuracy to measure agreement of classifier and behaviorist labeling on ground truth videos. For each bout found by either the labeler or classifier, an IoU frame overlap of 50% is considered a bout agreement. Two types of bout overlap are shown. Instance 1, the behaviorist labeled one continuous bout has start and end frames which are slightly off from the classifier’s bout. Instance 2, the behaviorist labeled a gap between bouts where the classifier labeled one continuous bout. C) Plot showing changes in performance with changing IoU thresholds. For the IoU of 0.50 (50%) bout-based precision, recall, and accuracy are 0.83, 0.763, and 0.871, respectively. D) Total licking time found by the behaviorist and classifier for each ground truth video (r=0.99). E) Total licking time across strain and dose for males and females found by the classifier. C57BL/6J had the highest licking response to formalin dose followed by C3H/HeJ, then BALB/cJ and A/J. F) Sample kinetic features of 5% formalin treated C57BL/6J mice are shown over time (time spent licking for 5 minute bins). G) C57BL/6J and C3H/HeJ have similar licking changes over time, with C57BL/6J showing the highest licking peakVal response to formalin, while BALB/cJ and A/J show very little peak activity. All error bars represent standard error of mean.

### 3.2 Features correlated with nociception

We first aimed to automate the scoring of behaviors in the open field formalin assay using JABS constructed behavioral classifiers, heuristic measures, and gait.JABS is an active learning system that uses 12 point pose coordinates and calculates distances and angles between points at each frame across a specified window to train a behavior classifier [40]. One behavior we classified is licking/biting. Bout overlap was used to determine success of this classifier, with 50% frame overlap between classifier and labeling behaviorist being considered a successfully labeled bout (Figure 1B). We achieved a precision of 0.91, a recall of 0.80, an accuracy of 0.90, and an F1 beta of 0.85 at bout-overlap threshold (IoU) of 0.50 (Figure 1C, Table **S2,S3**). Next, the time spent licking in each ground truth video was scored by classifier and behaviorist independently (Figure 1D). These measures were highly similar (r=0.99) (Figure 1D). Combined, these results show that automated and reliable scoring of licking is achieved.

We applied our validated classifier to the experimental dataset and observed effects of strain, dose, and sex on licking behavior (Figure 1 F). We also observed the previously reported bi-phasic licking response in this assay, with peaks at 5-10 and 15-35 minutes, supporting the accuracy of our trained classifier [29]. Some temporal aspects of licking behavior can be captured by kinetic features such as the time at the peak of licking behavior (peakTime), the value at the peak of licking behavior (peakVal), the value of the slope from start to licking peak (peakSlope), the length of time from the peak to where licking falls below 80% of the peak value (peakLength), and the value of the slope from the start to end of trial (timeSlope) (Figure 1F). These kinetic features show differences across dose and strain (Figure 1G). We found that C57BL/6J show higher licking, C3H/HeJ show a medium amount of licking, and BALB/cJ and A/J show less licking. Female C57BL/6J showing the most licking time response overall

We also trained a classifier for paw shaking, a known nocifensive behavior in the formalin assay where the mouse quickly shakes/flicks the injected paw. We find that time spent shaking increases with dose for all strains except C3H/HeJ (Figure 2). We next trained a classifier for rearing supported by a wall, as formalin has been shown to reduce rearing behavior [30]. Time spent rearing decreases with dose for C57BL/6J (Figure 2, Table **S4**).

**Figure 2:**
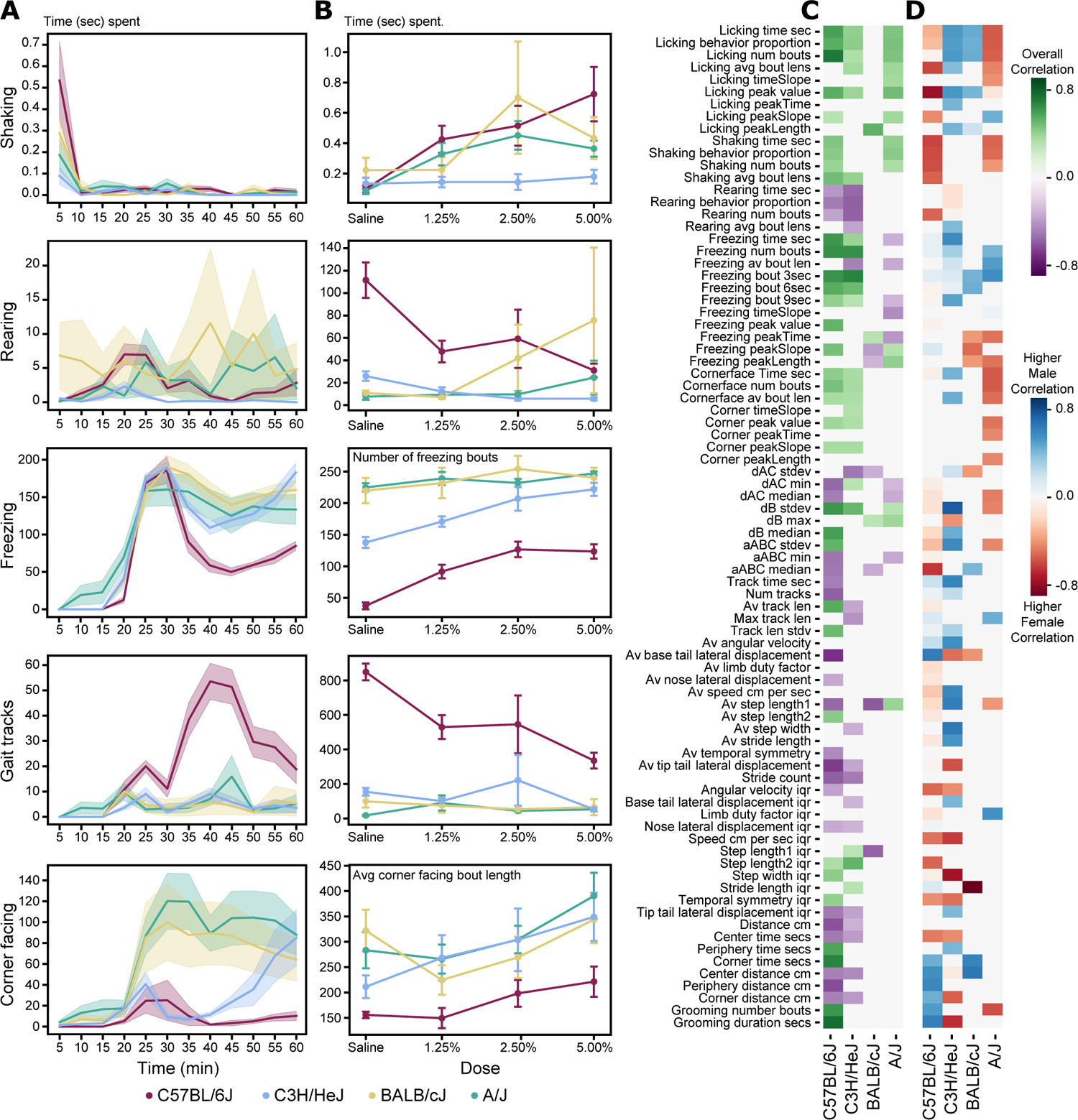
Behaviors correlated to formalin dose. A) Time spent in a behavior (shaking, rearing, freezing, gait, and corner facing) for mice injected with 5.00% formalin summed at each 5 minute bin for each strain. B) Summed sample features from each behavior across dose for each strain. C) Correlations between features and formalin dose for each strain. D) Sex differences in correlation between features and dose for each strain.

We observed that some mice given high doses of formalin had bouts of freezing (staying completely still for several seconds at a time) and corner facing (sitting completely still facing a corner), and assessed these behavioral heuristically. Number of freezing bouts increases with formalin dose for C57BL/6J and C3H/HeJ, but not for BALB/cJ and A/J which tend to have frequent freezing bouts even at saline doses (Figure 2, Table **S4**). The average length in seconds of a corner facing bout increased with dose for all strains except for BALB/cJ, which show long corner facing bout lengths at a saline dose (Figure 2A, Table **S4**).

We then considered gait using methods previously established by our group to extract stride measures from freely moving mice in the open field [36]. For many BALB/cJ and A/J mice, we found little to no strides throughout the video, resulting in missing data. We sought to quantify all gait tracks by calculating the speed of the center point of the mouse and finding frames above a threshold speed. Bout agreement accuracy was 0.93 and F1 beta was 0.84 for 20 minutes of densely labelled video for bouts greater than 10 frames (Table **S3**). Time spent in gait decreased with dose most clearly for C57BL/6J (Figure 2A, Table **S4**). When considering the bend of the spine we find significant correlation for these measures mostly for C57BL/6J, in particular dAC minimum and dB standard deviation (Figure 2C, Table **S4**).

In total, we analyzed 82 features (Table **S1**). For all classifiers and heuristics detailed, we achieved accuracies between 0.80 and 0.95 and F1 betas between 0.85 and 0.92 for the densely labeled video (Table **S3**). When we looked at the strengths of correlations across strains, we see an overall split between the higher activity strains (C57BL6J and C3H/HeJ) and the lower activity strains (A/J and BALB/cJ) (Figure 2C, Table **S4**). The higher activity strains showed a greater number of significant correlations to dose than the lower activity strains in gait features, location in the open field, freezing, and rearing. We find that the low activity strains tend to sit still near a corner for much of the open field test regardless of formalin dose. Thus, the nocifensive behaviors we score are less informative for these low activity strains.

### 3.3 Sex differences in nocifensive behavior

Sex differences in formalin response are often found for traditional features like licking but have not yet been established for some of our video features [28]. In order to investigate sex differences in our data by strain, we calculated the correlation of each feature with formalin dose for male and female mice in each strain. We then calculated the difference in correlation between males and females for each feature. For example, in C57BL/6J, males had a correlation between average distance from the corner and formalin dose of −0.62 while females had no significant correlation, resulting in a sex difference in correlation of 0.62 (Figure 2D). We found notable sex differences in formalin response in all 4 strains (Figure 2D). Licking behavior (ex: total licking time in seconds and the peak value of licking time) showed opposite strain specific sex differences; where C57BL/6J and A/J females had higher licking time, and in C3H/HeJ and BALB/cJ males were highest.

In C57BL/6J, the largest sex difference in formalin response was licking peak value, with a female dose correlation of 0.78 and no significant correlation for males. In this strain, features mostly related to licking, shaking, and gait (ex: time spent in gait, average temporal symmetry) showed higher response in females and features related to gait and locomotor changes (ex: average distance from corner, tip-tail lateral displacement IQR, step width IQR) showed higher responses for males. This suggests that while C57BL/6J females show stronger nociceptive response in traditional formalin measures like licking and shaking in addition to gait, males show more nociceptive response in gait features.

In C3H/HeJ, the largest sex difference in formalin response was average lateral displacement of the tail tip in stride with females having a correlation of −0.75 and males having no correlation. Features related to gait were more correlated for females (ex: tip-tail lateral displacement IQR, average step width) and features related to licking, freezing, engineered features, and gait (ex: max gait track length, number of gait tracks) had higher correlation for males. Interestingly, while both C57BL6J and C3H/HeJ are higher responders to formalin overall, the observed sex differences are in opposite directions.

For BALB/cJ, there was a large sex difference in formalin response was stride count, with females having a correlation of −0.95 with formalin dose and males having no significant correlation. However, it is important to note that only 5 of the 26 females had information for stride count. Features such as freezing peak slope and distance from the corner had higher correlations for females, and features related to licking, freezing bouts, and step length had higher correlation for males.

For A/J, the largest sex difference in formalin response was temporal symmetry IQR, with females having a correlation of 0.58 and males having no significant correlation. Features related to licking, shaking, freezing kinetics and corner facing having higher correlations for females, and features related mostly to freezing behavior and time spent in corner were higher for males.

### 3.4 Univariate Model of formalin dose

Univarite scales are useful for multiple scientific questions and can be constructed using different combinations of data from the 4 strains. For instance, studies using genetically diverse mice need a scale that generalizes across strains. Certain genetic and neurobiological studies are limited to specific strains, and a well constructed strain specific univariate model will be useful. Similarly, analgesic discovery can be carried out in specific strains, however one may choose to do preclinical work in genetically diverse populations that better model the human populations, necessitating several strain specific models. We modeled 7 univariate scales - 4 strain specific models, 1 model for *high* activity strains (C3H/HeJ and C57BL/6J), 1 model for *low* activity strains (A/J and Balb/cJ), 1 model for *all* strains. We hypothesized that the strain specific models would have the best performance. In addition, we tested if using additional features allows more accurate detection of nociception than the traditionally scored features of licking and shaking. To do this, we compared a univariate models using all our features and a model using just the traditional features. An idealized distribution of mice along the univariate scale is shown in Figure 3A. We see near complete separation between the mice of different doses along this ideal scale; each segment corresponding to a dose contains almost all of the mice injected with that dose.

**Figure 3:**
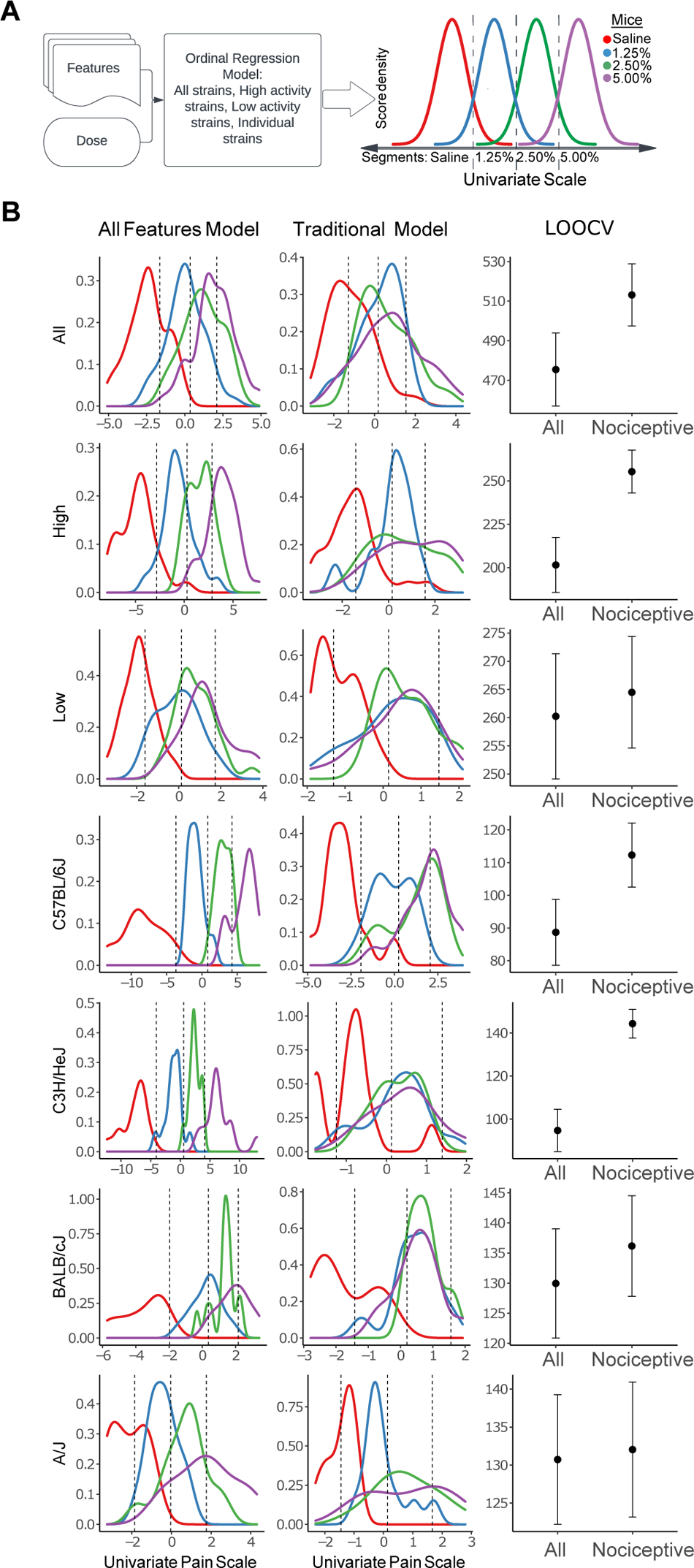
Creation of a univariate scale for dose detection. A) Outline of modeling process. Ordinal regression models were trained for 7 subsets of data (All strains, high activity strains, low activity strains, and the 4 individual strains). This resulted in a latent continuous scale with segments corresponding to each dose. The resulting distribution of mice along the scale shows an ideal separation of dose in each segment. B) Results for each model. The performance of the model using the expanded feature set (All features model) was compared to the performance of the model trained only using the 13 traditional formalin measures (licking and shaking).

We first looked at a global model for all strains. This model used all available features excluding gait features as many low activity strain (BALB/cJ and A/J) mice did not take enough strides to calculate these (Figure **S1**). We found that the univariate scale from the All Strain model using all video features more accurately discriminated between formalin doses (LOO=0.47) than the model using only the traditional nociception features (LOO=0.42; Figure 3B, Table 1). Some of the most important features for the model were freezing time, freezing average bout length, sex, and licking peak value (Figure **S2**).

**Table 1:** Leave one out (LOO) accuracy for each model

|  | Traditional nocifensive features |  |  | All features |  |  |
| --- | --- | --- | --- | --- | --- | --- |
|  | All | Male | Female | All | Male | Female |
| All Strains | 0.42 | 0.39 | 0.46 | 0.47 | 0.48 | 0.46 |
| High Activity | 0.45 | 0.44 | 0.46 | 0.56 | 0.54 | 0.58 |
| Low Activity | 0.34 | 0.31 | 0.37 | 0.37 | 0.35 | 0.39 |
| C57BL/6J | 0.5 | 0.5 | 0.5 | 0.6 | 0.69 | 0.5 |
| C3H/HeJ | 0.29 | 0.27 | 0.31 | 0.6 | 0.62 | 0.58 |
| BALB/cJ | 0.26 | 0.29 | 0.23 | 0.4 | 0.38 | 0.42 |
| A/J | 0.46 | 0.36 | 0.56 | 0.32 | 0.32 | 0.32 |

We hypothesized that model of high and low activity groups separately would increase accuracy. Indeed, for the High Activity model using all features, we achieved more accurate dose discrimination for the all feature model (LOO=0.56) compared to the traditional nociception model (LOO=0.45), and the global all strains model (LOO=0.47; Figure 3B, Table 1). Freezing measures like freezing time, freezing average bout length, and freezing time slope are among the most important features for the High Activity model but others include stride length IQR and average step length. For the Low activity strains model using all features except gait measures (59 features), we do not show significant improvement over the traditional nocifensive features model (Figure 3B, Table 1). This indicates, in our assays when strains do not show enough locomotor behavior, the traditional measures offer best performance.

Next we trained strain specfic models. Both the C57BL/6J and C3H/HeJ we achieved even higher dose discrimination accuracy (LOO=0.60 and 0.60, respectively; Figure 3B, Table 1). However, even though both strains are show high locomotor activity, the strain specific models use different features for discrimination. Thus,features which were important are markedly different between the C57BL/6J and C3H/HeJ models (Figure 4). For the C57BL/6J model, grooming duration was the most important feature by a large margin, followed by average step length, dAC minimum, and average step width. For the C3H/HeJ model, the most important features were average tip tail lateral displacement, grooming duration, licking bouts, and freezing time (Figure 4). The C57BL/6J model was also more accurate for males (LOO=0.69) than females (LOO=0.5). However, neither the BALB/cJ or A/J model achieved significantly higher accuracy than the traditional nociception model, although the BALB/cJ expanded feature model saw trending improvements over the traditional features model (Figure 3B, Table 1).

**Figure 4:**
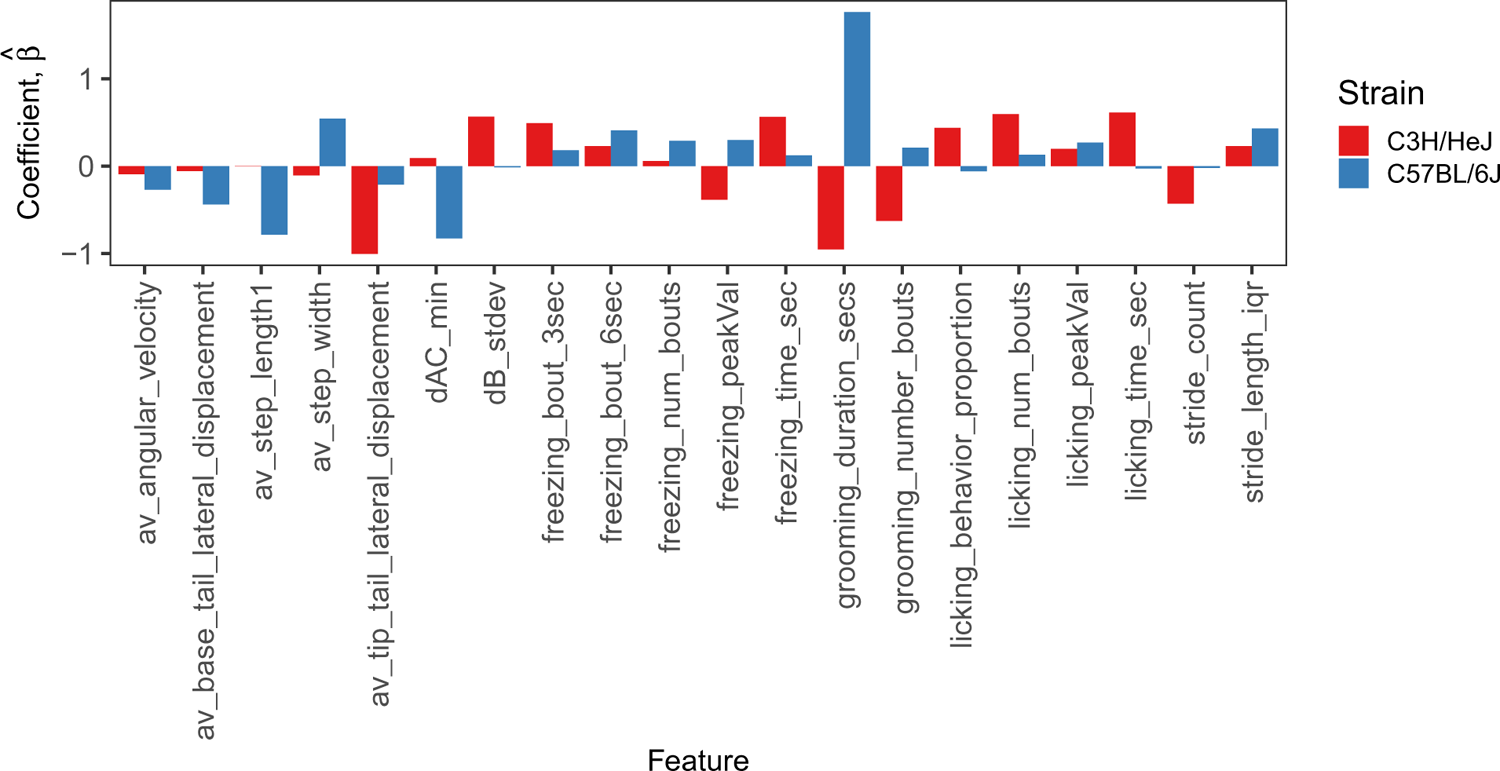
Top 10 important features for dose discrimination in C57BL/6J and C3H/HeJ specific models show differences. The heights of the bars represent the estimated posterior means, 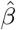.

### 3.5 Morphine induced antinociception

Given robust classification of formalin dose using our univariate model, we next assessed the ability of this model to detect morphine induced antinociception in the formalin assay. We expected to see the mice given no morphine with formalin to have high nociceptive scores while mice given morphine with formalin to have lower scores. Indeed, we find this overall pattern in our data (Figure 5). Although we do not see clear separation between 5 mg/kg and 10 mg/kg morphine administration in the univariate scale, there is clear separation between the two morphine groups and saline mice. We find that the mice given no morphine mostly fall along the 3rd and 4th segments of the univariate scale (corresponding to scores for 2.50% and 5.00% formalin) while both the mice given 5 mg/kg and 10 mg/kg morphine fall mostly within the second segment (corresponding to scores for 1.25% formalin). We used the center line on the univariate scale demarcating higher foramlin dose from lower formalin dose to test categorization accuracy: to the right of the line should be mice receiving no morphine and to the the left should be mice receiving morphine. We find an overall accuracy of 0.895, a male accuracy of 1.00, and a female accuracy of 0.792.

**Figure 5:**
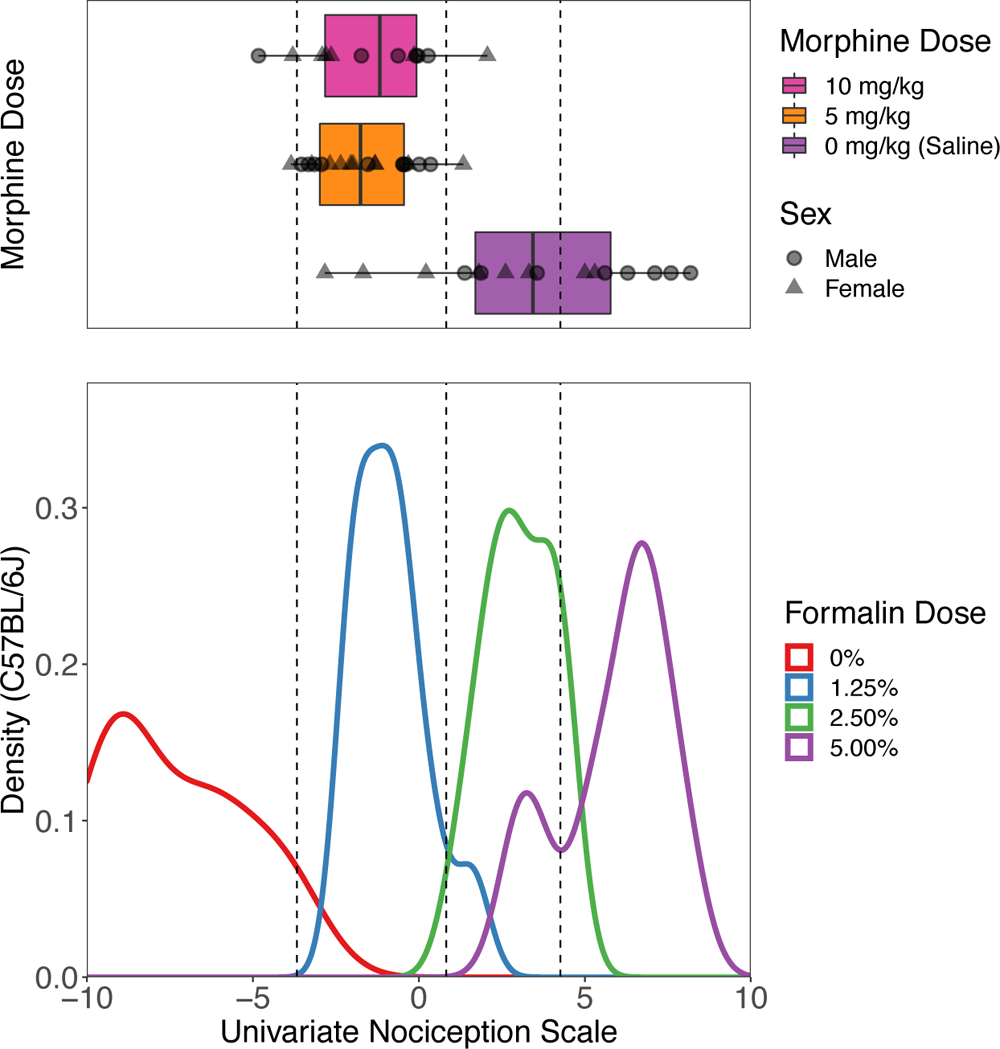
Model trained on C57BL/6J formalin data can detect antinociceptive effect of morphine. Mice given 5.00% formalin and various doses of morphine (0, 5, 10 mg/kg) were scored and their score along the univariate scale is shown. The mouse score distribution for the C57BL/6J model is shown below for comparison.

## 4 Discussion

We find that our machine vision approach has several advantages to scoring only traditional nocifensive features alone and addresses critical gaps in current experimental approaches to quantifying nociception. Our methods quantify voluntary evoked behaviors more naturalistic than operant and reflexive assays, in volumes impractical for manual scoring, at temporal and spatial resolutions not feasible for human observers, all at low cost. These advantages open opportunities to forward genetic studies requiring thousands of mice that were previously impractical, such as genetic mapping in diversity outbred panels. Because our univariate model samples 82 nocifensive responses, it allows for more appropriate comparison across strains with differing nocifensive behavioral structures. This allows increased confidence that observed differences are due to nociception and not confounding traits like anxiety, culminating in a pain assay better suited for wide genetics panels that will increase the validity of loci and genes identified through genetic mapping. Our approach also found a number of features associated with high formalin response previously unidentified as nocifensive behaviors, many demonstrating strain specific expression. This opens the possibility of interrogating the genetic origins of nociceptive behavioral structure to geneticists, which is not possible with traditional metrics alone.

Sex differences in rodent nociception have been a topic of increased interest in recent years, and females have historically been understudied in biomedical research. The emerging literature points to fundamental sex differences in nociception and analgesia, and significant interactions between strain and sex [14, 15, 41–44]. We identified sex differences in all 4 strains, many of which are strain specific. For example, while the high activity strains C57BL/6J and C3H/HeJ are overall high responders to formalin, they have opposing sex differences in response to licking and other features. We also observed higher nociceptive responses in female C57BL/6J and A/J, while C3H/HeJ and BALB/cJ showed higher responses in males. Finally, in our “All Strain” formalin dose prediction model, sex was one of the most important features. This supports observed interactions between strain, sex, and nociception, but also suggests that tests optimized to assay nociception in male mice may lack sensitivity in females and continue to under serve this population in biomedical research. We think our machine learning approach offers a more equitable and objective tool to study pain in female subjects and the interaction between sex and genotype.

There is a dire need for novel therapeutic interventions for pain. Current pain interventions rely primarily on opioids, which are accompanied by dangerous clinical complications. Using our C57BL6/J strain specific model, we observed robust discrimination between morphine and control mice. This modeling was most successful in males compared to females. In C57BL/6J, males have been shown to be significantly more sensitive to morphine compared to females which may account for the higher behavioral correlations in males [45]. Given their ability to discern the antinociceptive effects of the analgesic morphine, this approach open the door to high throughput screening for novel analgesic therapeutics. By sampling more naturalistic responses to noxious stimuli these methods also bring us closer to the human condition, potentially improving the translatability of pre-clinical work to human intervention. As these methods can also build robust indices for pain across strains, they offer a tool to interrogate antinociception on discrete genetic backgrounds, further increasing the tranlatability of pre-clinical work.

These data demonstrate the success of machine vision-based univariate scales in acute pain models, but could be scaled for multi-day assessment of nociception in more complex environments. Such advancements would facilitate the study of the progression and treatment of chronic pain by using more naturalistic behaviors, potentially improving translatability of biological and pharmacological findings. Furthermore, prolonged behavioral testing could incorporate quantification of disrupted eating, social interaction, and sleep, outcomes observed in mouse models of pain [46–48]. These features would improve the assessment of both chronic and acute pain models and could be discriminatory features to improve quantification of pain in low activity strains.

Overall, this machine vision platform provides a complimentary approach to existing machine learning approaches that assess pain, such as facial grimace and high-speed videography, by allowing free mobility of each animal and reducing the overall data burden needed to asses pain phenotypes over prolonged periods [49–52]. However, it possible these approaches could improve quantification of nociception in mice with low overall activity. Our univariate scale is a high-throughput, sensitive, and low cost tool suited for researchers studying the genetics of and pharmacological interventions for nociception. Its ability to quantify nociception across strains and sexes makes it possible to compare and contrast disparate strains and sexes and facilitates the direct comparison between mice with differing nociceptive behavioral structures.

## 5 Methods

### 5.1 Mice

C57BL/6J (000664), C3H/HeJ (000659), BALB/cJ (000651), and A/J (000646) mice were obtained from the Jackson Laboratory. Mice were between 2-3 months old at time of testing. Mice were single sex, group-housed (3–5) with ad lib water and food under a 12-hour light–dark schedule (0600-1800), and experiments were conducted in the light phase. All procedures and protocols were approved by The Jackson Laboratory Animal Care and Use Committee and were conducted in compliance with Institutional Animal Care and Use Committee.

### 5.2 Open field assay

The open field behavioral assays were conducted as previously described using a top down camera view [39, 53]. Prior to testing, mice were allowed to acclimate to the testing room for 30–45 minutes. Mice were habituated to the open field or 15 minutes. They were then anesthetised with isoflurane 4% (Henry Schein Isothesia; 1169567762) and the left hind paw of the mouse was injected (intra-plantar) with 30 μl of 0%, 1.25%, 2.50%, or 5.00% formalin (formaldehyde solution, Sigma-Aldrich; 15512) in saline (sterile saline, Henry Schein, 002477). Mice were placed back in the open field arena for recovery and recorded for another 60 minutes. Mice were euthanized the same day they were injected with formalin. Each dosage was tested on at least 5 males and 5 females for each strain, resulting in a dataset of 194 mice.

### 5.3 Video recording, segmentation, and tracking

Our open field arena, video apparatus, and tracking and segmentation networks are as detailed previously [39]. Briefly, the open field arena measures 20.5” by 20.5” with Sentech camera mounted 40 inches above. The camera collects data at 30 fps with a 640×480px resolution. We use a neural network trained to produce a segmentation mask of the mouse to produce an ellipse fit of the mouse at each frame as well as a mouse track.

### 5.4 Pose estimation and gait

The 12-point 2D pose estimation was produced using a deep convolutional neural network trained as previously detailed [24]. The points captured are nose, left ear, right ear, base of neck, left forepaw, right forepaw, mid spine, left rear paw, right rear paw, base of tail, mid tail and tip of tail. Each point at each frame has an x coordinate, a y coordinate, and a confidence score. We use a minimum confidence score of 0.3 to determine which points are included in the analysis.

The gait metrics were produced as detailed in Shephard et al. (2022) [24]. Briefly, the stride cycles were defined by starting and ending with the left hind paw strike, tracked by the pose estimation. These strides were then analyzed for several temporal, spatial, and whole-body coordination characteristics, producing gait metrics. Our gait method is also conservative with the strides it collects; briefly, it only includes high confidence strides and it discards the first and last stride of each gait track, thus requiring a gait track to be at least 3 strides long [36]. Each stride cycle was analyzed for its spatial, temporal, and postural measures (ex: step width, nose-lateral displacement). We took the averages and inter-quartile ranges for all measures. We previously developed methods for looking at the bend of the spine throughout the video [54]. Briefly, we track the pose coordinates of three points on the mouse at each frame: the back of the head (A), the middle of the back (B), and the base of the tail (C).

### 5.5 JABS classifiers

Full detail can be found in Beane et al. (2022) [40]. Briefly, our data acquisition uses custom designed standardized data acquisition hardware and software that provides a controlled environment, optimized video storage, and live monitoring capabilities. This is leveraged by the JABS annotation and classification software which is a python based GUI utility for behavior annotation and training classifiers using the annotated data. One can then use the trained classifiers for predicting whether behavior happens or not in the unlabeled frames. For this study behavioral classifiers were made to assess licking, paw shaking, and rearing. While machine learning techniques are often evaluated using frame accuracies, behaviorists find detecting the same bouts of behavior more important than the exact starting and ending frame of these bouts. Even between two humans labeling the same behavior, there are unavoidable discrepancies in the exact frames of the behavior. Therefore, we use a bout-based accuracy rather than a frame-based accuracy to evaluate the performance of the classifier. For each bout found by either the classifier or the behaviorist, the frames within that bout where both the classifier and behaviorist agree was calculated. If at least half of the frames agreed, we consider this bout to be overlapping. The licking classifier was trained on a set of videos from a prior experiment using male and female C57BL/6J mice. A behaviorist densely labeled (i.e., labeled each frame of video as behavior or not-behavior) 6 videos from each of the 4 strains of mice injected with 5.00% formalin for a total of 1 hour of ground truth labelling. Trained classifiers were assessed for accuracy by comparing using “ground truth videos,” where classifier and behaviorist independently graded behavior, and results were compared.

### 5.6 Open field measures and feature engineering

Open field measures were derived from ellipse tracking of mice as described before [35, 39]. The tracking was used to produce locomotor activity and anxiety features. Grooming was classified using a action detection network as previously described [35]. Spinal mobility metrics and rearing were derived using the pose estimation data as previously described [54]. Freezing behavior was heuristically derived by taking the average speed of the nose, base of head, and base of tail points at each frame, and finding periods of at least 3 seconds where the average speed of the mouse was less than 0.09 pixels/sec. Corner-facing behavior was heuristically derived by taking the intersection of frames where freezing behavior occurred and frames where the nose point was within 25 pixels of an arena corner.

### 5.7 Validations

For each bout of behavior found by either the classifier/heuristic, the frames within that bout where both the classifier and a behaviorist agree that the behavior occurs was calculated. If at least 50% of the frames agreed, we considered this bout to be a true positive. This was repeated for each bout of “not-behavior”, and overlapping bouts of non-behavior were considered true negatives. If less than 50% of the classifier/heuristic bout was in agreement with the behaviorist, it was considered a false positive or a false negative.

### 5.8 Behavioral correlation with dose

We used Pearson’s correlation measure to compute associations between formalin dose and behavioral outcomes. For example, we extracted the two variables dose and licking time (secs) for C57BL/6J mice to estimate the correlation. To determine the difference in correlation between sex within strains, male and female correlations were calculated and female Pearson’s R was subtracted from the male value, with no significant correlation treated as a value of 0.

### 5.9 Modeling

Here we describe the univariate response model for formalin dose. We used ordinal regression modeling to produce a univariate scale: a continuous latent variable which underlies the formalin doses. Each mouse lies somewhere along this scale based on its feature values. We used the linear combination of input features for prediction via the threshold parameters to divide the continuous univariate scale into segments. Each segment corresponds to one of the formalin dosages. We created dose predicting models using all mice, but excluded gait features due to low representation in low activity strains. We also created models specific for high activity strains, low activity strains, and each of the 4 strains individually.

We briefly describe cumulative link models, which are the most common regression models for analyzing ordinal data [55]. These models assume that the categories of the observed response variable, Y, can be modeled as contiguous intervals on a continuous latent scale (i.e., not observable) *Y*\*. More formally, the model can be written as follows:

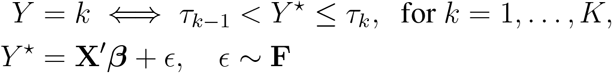

where *Y*, the subject’s response (Dose), is one of the *K* ordinal categories (K=4; 0%, 1.25%, 2.50%, 5.00%), **X**= (*X*_1_,…, *X_p_*}′ denotes a vector of subject’s observed *p* covariates (features), *β* = (*β*_1_,…,*β*_p_) denotes the vector of unknown regression coefficients to be inferred. Assuming F to be a standard logistic distribution, the cumulative *logit* link model has the form,

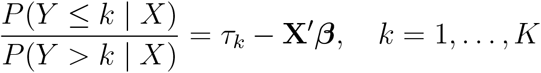

where the intercepts *τ_k_* satisfy the constraint —∞= *τ*_0_ < *τ*_1_ <… < *τ_K_* < *τ_K_*_+1_ = ∞. The *τ*’s are estimated from the data and are shown as the vertical dotted/dashed lines in Figures 3B, 5.

We adopt a Bayesian approach [56] and place a regularized horseshoe shrinkage prior distribution [57, 58] on the model parameters to address multi-collinearity and identify relevant features. The regularized horseshoe is defined as follows.

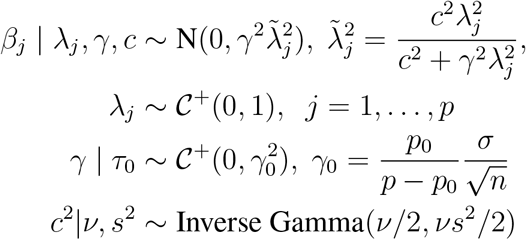

where *p*_0_ denotes the prior guess on the number of relevant features, *N*, 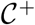 are univariate Gaussian and half-Cauchy distributions respectively. Following the recommendations of [59], we set *v* = 4, *s*^2^ = 2. Since we do not have apriori information on the number of relevant variables, we used a half-Cauchy prior on the global shrinkage parameter *γ*, i.e. 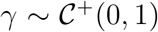. We placed a uniform prior for *τ* subject to the constraint *τ*_1_ < *τ*_2_ <… < *τ_K_*[60].

We standardized the features by subtracting the mean and dividing by the standard deviation of the feature. We fit our models using four randomly initialized chains, each with 2000 iterations, of which the first 1000 were warm-up iterations to calibrate the No-U-Turn-Sampler (NUTS) [61], leading to a total of 4000 posterior samples. We evaluated the convergence of the sampler using the split 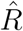 statistic [62]. The 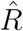 value provides information on the convergence of the algorithm. We calculated 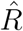 and found that none of the chains had 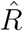 considerably greater than 1 (i.e., > 1.1) establishing that all chains had converged. We performed posterior computations using Stan [63].

### 5.10 Assessment of morphine induced antinociception

We next used a morphine experiment to test if our model could distinguish between mice given morphine and mice not given morphine during formalin testing. We administered one of 3 doses of morphine in 0.9We used the center line on the univariate scale demarcating higher foramlin dose from lower formalin dose to test categorization accuracy: to the right of the line should be mice receiving no morphine and to the the left should be mice receiving morphine.

## 6 Acknowledgements

We thank Sean Deats, Tionna Ouellette, Brian Geuther, Keith Sheppard, and other members of the Kumar Lab for helpful advice and technical assistance. We thank JAX Information Technology team members Edwardo Zaborowski, Shane Sanders, Rich Brey, David McKenzie, and Jason Macklin for infrastructure support. This work was funded by The Jackson Laboratory Directors Innovation Fund, National Institute of Health DA041668 (NIDA), DA048634 (NIDA). All code and training data will be available at Kumarlab.org and Kumar Lab Github.

