## Supplemental Tables and Figures for "A high-throughput machine vision-based univariate scale for pain and analgesia in mice"

December 29, 2022

### 1 Supplement

Table S1: Video Features

| Video Features | | | |
|---|---|---|---|
| Category | Name | Description | Units |
| | <behavior>_time_sec | The amount of time the mouse spent in behavior | sec |
| | <behavior>_behavior _proportion | The proportion of behavior time to total video time | sec |
| | <behavior>_num_bouts | The number of behavior bouts for the mouse. | $\sim$ |
| | <behavior>_avg _bout_lens | The average length of time for a behavior bout. | sec |
| | <behavior>_peakTime | the time at the peak of licking behavior. | $\sim$ |
| | <behavior>_peakVal | Across 5 minute bins, the value at the peak of behavior. | $\sim$ |
| | <behavior>_peakSlope | Across 5 minute bins, the value of the slope from start to behavior peak. | $\sim$ |
| | <behavior>_peakLength | Across 5 minute bins, the length of time from the peak to where behavior falls below 80% of the peak value. | $\sim$ |
| | <behavior>_timeSlope | Across 5 minute bins, the value of the slope from the start to end of trial. | $\sim$ |
| Traditional | licking | Behavior where the mouse licks/bites the injected hind paw. | $\sim$ |
| Traditional | shaking | Behavior where mouse quickly shakes the injected paw. | $\sim$ |
| Engineered | rearing | Behavior where the mouse stands on its hind legs while supported by an arena wall. | $\sim$ |
| Engineered | freeze | Behavior where the mouse sits in complete stillness for >3 seconds. | $\sim$ |

| Video Features | | | |
|---|---|---|---|
| Category | Name | Description | Units |
| Engineered | freeze_bout_3sec | The number of freezing bouts between 3 and 6 seconds. | ∼ |
| Engineered | freeze_bout_6sec | The number of freezing bouts between 6 and 9 seconds. | ∼ |
| Engineered | freeze_bout_9sec | The number of freezing bouts greater than 9 seconds. | ∼ |
| Engineered | cornerface | Behavior where the mouse is freezing while its nose is facing an arena corner. | ∼ |
| Engineered | num_tracks | The number of bouts of continuos gait for the mouse. | ∼ |
| Engineered | max_track_len | The longest bout of a gait track. | sec |
| Engineered | track_len_stdv | The standard deviation of the length in time of all bouts of a gait track. | sec |
| Engineered | av_track_len | The average length in time of a gait track. | sec |
| Open Field | distance_cm | Sum of locomotor activity. | cm |
| Open Field | center_time_secs | Sum of time spent in center. | sec |
| Open Field | periphery_time_secs | Sum of time spent along any wall. | sec |
| Open Field | corner_time_secs | Sum of time spent in any corner. | sec |
| Open Field | center_distance_cm | Average distance from center across the video. | cm |
| Open Field | periphery_distance_cm | Average distance from nearest periphery across the video. | cm |
| Open Field | corner_distance_cm | Average distance from nearest corners across the video. | cm |
| Open Field | grooming_number_bouts | Sum of all grooming bouts in video. | ∼ |
| Open Field | grooming_duration_secs | Average length of grooming bouts. | sec |
| Gait | median_angular_velocity | The first derivative of angle of a mouse, determined by the vector connecting the mouse's base of tail to its base of neck | deg/sec |
| Gait | median_base_tail _lateral_displacement | The difference between the minimum and maximum values of the base of tail's perpendicular distance from the mouse's displacement vector for a stride for each frame of a stride. Normalized by the mouse's body length. | ∼ |
| Gait | median_limb _duty_factor | The amount of time that the paw is in contact with the ground divided by the full stride time, calculated and averaged for each hind paw. | ∼ |
| Gait | median_nose _lateral_displacement | Calculated the same as base_tail_lateral_displacement, but using the nose point. | ∼ |
| Gait | median_speed_cm _per_sec | Speed is determined by the base of tail point. | cm/sec |

| Video Features | | | |
|---|---|---|---|
| Category | Name | Description | Units |
| Gait | median_step_length1 | The distance that the right hind paw travels past the previous left hind paw strike. | cm |
| Gait | median_step_length2 | The distance that the left hind paw travels past the previous right hind paw strike. | cm |
| Gait | median_step_width | The length of the shortest line segment that connects the right hind paw strike to the line that connects the left hind paw's toe-off location to its subsequent foot strike position. | cm |
| Gait | median_stride_length | The full distance that the left hind paw travels for a stride, from toe-off to foot-strike. | cm |
| Gait | median_temporal _symmetry | The difference in time between the left and right hindpaw strike, divided by the total strike time. | ~ |
| Gait | median_tip_tail _lateral_displacement | Calculated the same as base_tail_lateral_displacement, but using the tip-tail point. | ~ |
| Gait | stride_count | Sum of all recorded strides in video. | ~ |
| Gait | distance_cm_sc | Sum of locomotor activity, normalized by time spent in open field. | cm/sec |
| End of Table | | | |

Table S2: Ground truth validation confusion matrix

| | True Positive | False Positive | True Negative | False Negative |
|---|---|---|---|---|
| Licking | 32 | 3 | 71 | 8 |
| Shaking | 45 | 11 | 61 | 5 |
| Rearing | 37 | 2 | 49 | 5 |
| Freezing | 98 | 8 | 58 | 8 |
| Corner facing | 18 | 4 | 6 | 0 |
| Gait track | 109 | 8 | 137 | 34 |

Table S3: Ground truth validation accuracy

| | Precision | Recall | Accuracy | F1beta |
|---|---|---|---|---|
| Licking | 0.914 | 0.800 | 0.904 | 0.853 |
| Shaking | 0.803 | 0.900 | 0.868 | 0.849 |
| Rearing | 0.948 | 0.880 | 0.924 | 0.913 |
| Freezing | 0.924 | 0.924 | 0.906 | 0.924 |
| Corner facing | 0.818 | 1.000 | 0.857 | 0.900 |
| Gait track | 0.931 | 0.762 | 0.854 | 0.838 |

Table S4: Feature correlations (Pearson) with dose by strain

| Feature correlation with dose. | | | | | | | | |
|---|---|---|---|---|---|---|---|---|
| Features | C57BL/6J | | C3H/HeJ | | BALB/cJ | | AJ | |
| | Corr | $p_{value}$ | Corr | $p_{value}$ | Corr | $p_{value}$ | Corr | $p_{value}$ |
| licking_time_sec | 0.588 | 0 | 0.403 | 0.003 | 0.273 | 0.057 | 0.45 | 0.001 |
| licking_behavior _proportion | 0.537 | 0 | 0.418 | 0.002 | 0.263 | 0.068 | 0.462 | 0.001 |
| licking_num_bouts | 0.716 | 0 | 0.324 | 0.019 | 0.229 | 0.114 | 0.455 | 0.001 |
| licking_avg _bout_lens | 0.157 | 0.272 | 0.364 | 0.008 | 0.222 | 0.147 | 0.378 | 0.008 |
| licking_timeSlope | 0.265 | 0.06 | 0.093 | 0.511 | 0.079 | 0.592 | 0.364 | 0.009 |
| licking_peakVal | 0.535 | 0 | 0.385 | 0.005 | 0.247 | 0.087 | 0.478 | 0 |
| licking_peakTime | -0.072 | 0.618 | 0.111 | 0.433 | -0.193 | 0.184 | -0.085 | 0.555 |
| licking_peakSlope | 0.301 | 0.032 | 0.23 | 0.101 | 0.19 | 0.19 | 0.367 | 0.009 |
| licking_peakLength | 0.072 | 0.618 | -0.111 | 0.433 | 0.524 | 0 | 0.033 | 0.822 |
| shaking_time_sec | 0.45 | 0.001 | 0.102 | 0.471 | 0.162 | 0.265 | 0.406 | 0.004 |
| shaking_behavior _proportion | 0.429 | 0.002 | 0.078 | 0.582 | 0.156 | 0.284 | 0.429 | 0.002 |
| shaking_num_bouts | 0.387 | 0.005 | 0.007 | 0.963 | 0.119 | 0.415 | 0.338 | 0.016 |
| shaking_avg _bout_lens | 0.48 | 0.002 | 0.395 | 0.046 | 0.158 | 0.395 | 0.155 | 0.354 |
| rearing_time_sec | -0.407 | 0.003 | -0.562 | 0 | 0.198 | 0.174 | 0.204 | 0.155 |
| rearing_behavior _proportion | -0.494 | 0 | -0.577 | 0 | 0.195 | 0.179 | 0.205 | 0.152 |
| rearing_num_bouts | -0.362 | 0.009 | -0.571 | 0 | 0.203 | 0.162 | 0.224 | 0.119 |
| rearing_avg _bout_lens | 0.036 | 0.803 | -0.362 | 0.009 | -0.159 | 0.275 | -0.167 | 0.25 |
| freezing_time_sec | 0.615 | 0 | 0.39 | 0.006 | 0.018 | 0.902 | -0.318 | 0.028 |
| freezing_num_bouts | 0.639 | 0 | 0.636 | 0 | 0.136 | 0.353 | 0.096 | 0.516 |
| freezing_av _bout_len | -0.05 | 0.734 | -0.482 | 0.001 | -0.186 | 0.202 | -0.345 | 0.017 |
| freezing_bout_3sec | 0.631 | 0 | 0.675 | 0 | 0.205 | 0.157 | 0.244 | 0.095 |
| freezing_bout_6sec | 0.545 | 0 | 0.495 | 0 | 0.192 | 0.186 | 0.061 | 0.683 |
| freezing_bout_9sec | 0.404 | 0.004 | 0.299 | 0.039 | -0.1 | 0.496 | -0.296 | 0.041 |
| freezing_timeSlope | 0.262 | 0.061 | -0.033 | 0.818 | 0.131 | 0.364 | -0.438 | 0.002 |
| freezing_peakVal | 0.515 | 0 | -0.101 | 0.475 | -0.184 | 0.201 | -0.278 | 0.05 |
| freezing_peakTime | -0.016 | 0.909 | -0.19 | 0.178 | 0.3 | 0.034 | -0.402 | 0.004 |
| freezing_peakSlope | 0.466 | 0.001 | 0.208 | 0.138 | -0.385 | 0.006 | 0.303 | 0.032 |
| freezing_peakLength | 0.016 | 0.909 | 0.19 | 0.178 | -0.3 | 0.034 | 0.402 | 0.004 |
| cornerface_time_sec | 0.419 | 0.002 | 0.348 | 0.012 | -0.063 | 0.662 | 0.274 | 0.055 |
| cornerface_num_bouts | 0.465 | 0.001 | 0.352 | 0.011 | -0.09 | 0.534 | 0.247 | 0.084 |
| cornerface_av _bout_len | 0.345 | 0.031 | 0.333 | 0.021 | 0.091 | 0.531 | 0.138 | 0.34 |
| cornerface_timeSlope | 0.208 | 0.138 | 0.315 | 0.024 | 0.179 | 0.212 | 0.094 | 0.517 |

| Feature correlation with dose. (Table continued) | | | | | | | | |
|---|---|---|---|---|---|---|---|---|
| Features | C57BL/6J | | C3H/HeJ | | BALB/cJ | | AJ | |
| | Corr | $p_{value}$ | Corr | $p_{value}$ | Corr | $p_{value}$ | Corr | $p_{value}$ |
| cornerface_peakVal | 0.379 | 0.006 | 0.334 | 0.017 | -0.089 | 0.539 | 0.258 | 0.07 |
| cornerface_peakTime | 0.263 | 0.06 | 0.03 | 0.837 | 0.071 | 0.624 | 0.17 | 0.237 |
| corner_peakSlope | 0.352 | 0.011 | 0.346 | 0.013 | -0.104 | 0.471 | 0.078 | 0.592 |
| cornerface_peakLength | 0.132 | 0.35 | -0.124 | 0.387 | -0.069 | 0.635 | -0.17 | 0.237 |
| dAC_stdev | -0.082 | 0.564 | -0.513 | 0 | -0.317 | 0.025 | -0.11 | 0.449 |
| dAC_min | -0.558 | 0 | 0.3 | 0.032 | -0.137 | 0.341 | -0.287 | 0.043 |
| dAC_median | -0.487 | 0 | 0.108 | 0.45 | -0.098 | 0.498 | -0.332 | 0.019 |
| dB_stdev | 0.625 | 0 | 0.503 | 0 | 0.25 | 0.08 | 0.3 | 0.034 |
| dB_max | 0.03 | 0.84 | -0.038 | 0.793 | 0.287 | 0.043 | 0.379 | 0.007 |
| dB_median | 0.594 | 0 | 0.197 | 0.167 | 0.278 | 0.051 | -0.127 | 0.38 |
| aABC_stdev | 0.517 | 0 | 0.217 | 0.127 | 0.207 | 0.15 | 0.269 | 0.059 |
| aABC_min | -0.518 | 0 | 0.147 | 0.303 | -0.152 | 0.292 | -0.351 | 0.013 |
| aABC_median | -0.51 | 0 | -0.122 | 0.393 | -0.335 | 0.017 | -0.042 | 0.775 |
| distance_cm | -0.618 | 0 | -0.312 | 0.024 | -0.234 | 0.105 | 0.242 | 0.091 |
| center_time_secs | -0.505 | 0 | -0.394 | 0.004 | -0.093 | 0.525 | -0.098 | 0.501 |
| periphery_time_secs | 0.574 | 0 | 0.216 | 0.124 | 0.074 | 0.613 | 0.053 | 0.716 |
| corner_time_secs | 0.676 | 0 | 0.041 | 0.774 | -0.121 | 0.408 | -0.22 | 0.125 |
| center_distance_cm | -0.48 | 0.001 | -0.447 | 0.001 | -0.082 | 0.576 | 0.067 | 0.646 |
| periphery_distance_cm | -0.627 | 0 | -0.269 | 0.053 | -0.24 | 0.096 | 0.273 | 0.056 |
| corner_distance_cm | -0.483 | 0 | -0.395 | 0.004 | -0.264 | 0.067 | -0.035 | 0.812 |
| grooming_number _bouts | 0.623 | 0 | -0.194 | 0.168 | -0.107 | 0.466 | 0.254 | 0.075 |
| grooming_duration _secs | 0.751 | 0 | -0.158 | 0.265 | -0.06 | 0.683 | 0.27 | 0.058 |
| track_time_sec | -0.485 | 0.001 | -0.112 | 0.45 | -0.109 | 0.456 | 0.156 | 0.29 |
| num_tracks | -0.578 | 0 | -0.149 | 0.313 | -0.168 | 0.248 | 0.17 | 0.248 |
| av_track_len | 0.548 | 0 | -0.353 | 0.014 | -0.18 | 0.236 | 0.026 | 0.87 |
| max_track_len | -0.023 | 0.875 | -0.416 | 0.003 | -0.132 | 0.388 | 0.113 | 0.476 |
| track_len_stdv | 0.499 | 0 | -0.283 | 0.051 | -0.09 | 0.558 | 0.144 | 0.363 |
| av_angular_velocity | -0.256 | 0.067 | -0.121 | 0.401 | -0.35 | 0.201 | 0.041 | 0.835 |
| av_base_tail _lateral_displacement | -0.698 | 0 | -0.183 | 0.203 | 0.165 | 0.558 | -0.028 | 0.89 |
| av_limb_duty_factor | 0.207 | 0.141 | 0.185 | 0.198 | 0.224 | 0.422 | -0.01 | 0.959 |
| av_nose _lateral_displacement | -0.347 | 0.012 | -0.185 | 0.199 | 0.17 | 0.546 | -0.061 | 0.76 |
| av_speed_cm_per_sec | -0.229 | 0.102 | -0.198 | 0.167 | -0.306 | 0.268 | 0.029 | 0.886 |
| av_step_length1 | -0.562 | 0 | -0.234 | 0.102 | -0.591 | 0.02 | 0.391 | 0.04 |
| av_step_length2 | 0.424 | 0.002 | 0.03 | 0.837 | 0.014 | 0.96 | 0.031 | 0.876 |
| av_step_width | -0.145 | 0.304 | -0.322 | 0.023 | -0.078 | 0.782 | -0.147 | 0.457 |
| av_stride_length | -0.178 | 0.207 | -0.257 | 0.072 | -0.465 | 0.08 | -0.05 | 0.801 |
| av_temporal_symmetry | -0.449 | 0.001 | -0.185 | 0.203 | -0.294 | 0.308 | -0.039 | 0.845 |

| Feature correlation with dose. (Table continued) | | | | | | | | |
|---|---|---|---|---|---|---|---|---|
| Features | C57BL/6J | | C3H/HeJ | | BALB/cJ | | AJ | |
| | Corr | $p_{value}$ | Corr | $p_{value}$ | Corr | $p_{value}$ | Corr | $p_{value}$ |
| av_tip_tail _lateral_displacement | -0.672 | 0 | -0.357 | 0.011 | 0.427 | 0.113 | -0.049 | 0.804 |
| stride_count | -0.58 | 0 | -0.499 | 0 | -0.222 | 0.361 | 0.25 | 0.125 |
| angular_velocity_iqr | -0.368 | 0.007 | -0.111 | 0.441 | 0.463 | 0.082 | 0.266 | 0.172 |
| basetail_lateral _displacement_iqr | -0.248 | 0.076 | -0.305 | 0.031 | 0.281 | 0.311 | 0.134 | 0.498 |
| limb_duty_factor_iqr | -0.071 | 0.617 | -0.114 | 0.429 | -0.391 | 0.15 | 0.081 | 0.683 |
| nose_lateral _displacement_iqr | -0.344 | 0.013 | -0.309 | 0.029 | -0.101 | 0.721 | -0.114 | 0.565 |
| speed_cm_per_sec _iqr | -0.214 | 0.128 | -0.232 | 0.105 | -0.375 | 0.169 | 0.293 | 0.13 |
| step_length1_iqr | 0.179 | 0.205 | 0.283 | 0.046 | -0.563 | 0.029 | 0.039 | 0.842 |
| step_length2_iqr | 0.336 | 0.015 | 0.518 | 0 | -0.404 | 0.136 | 0.116 | 0.557 |
| step_width_iqr | 0.418 | 0.002 | -0.206 | 0.151 | 0.421 | 0.118 | 0.042 | 0.831 |
| stride_length_iqr | 0.149 | 0.292 | 0.299 | 0.035 | -0.171 | 0.542 | 0.242 | 0.215 |
| temporal_symmetry _iqr | 0.401 | 0.004 | 0.032 | 0.829 | -0.155 | 0.598 | 0.124 | 0.531 |
| tip_tail_lateral _displacement_iqr | -0.502 | 0 | -0.369 | 0.008 | 0.08 | 0.778 | 0.038 | 0.849 |
| End of Table | | | | | | | | |

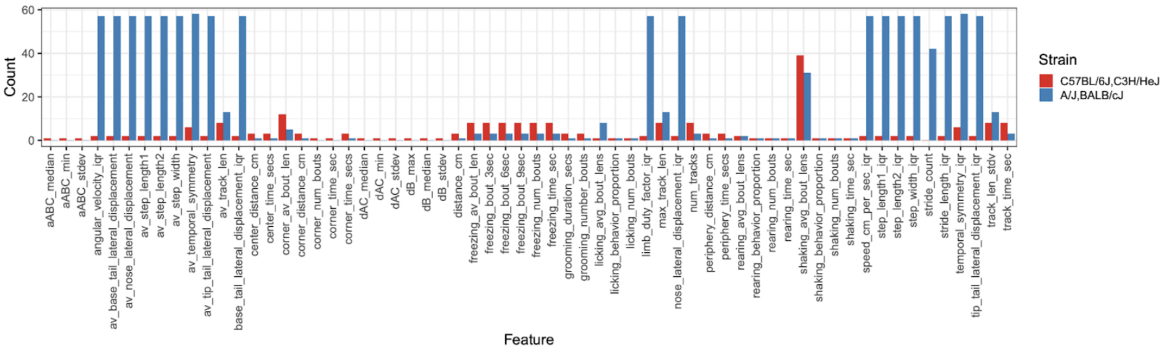


Figure S1: Low activity strains have many mice with missing gait feature values. The amount of missing data for each feature for high vs low activity mice is shown.

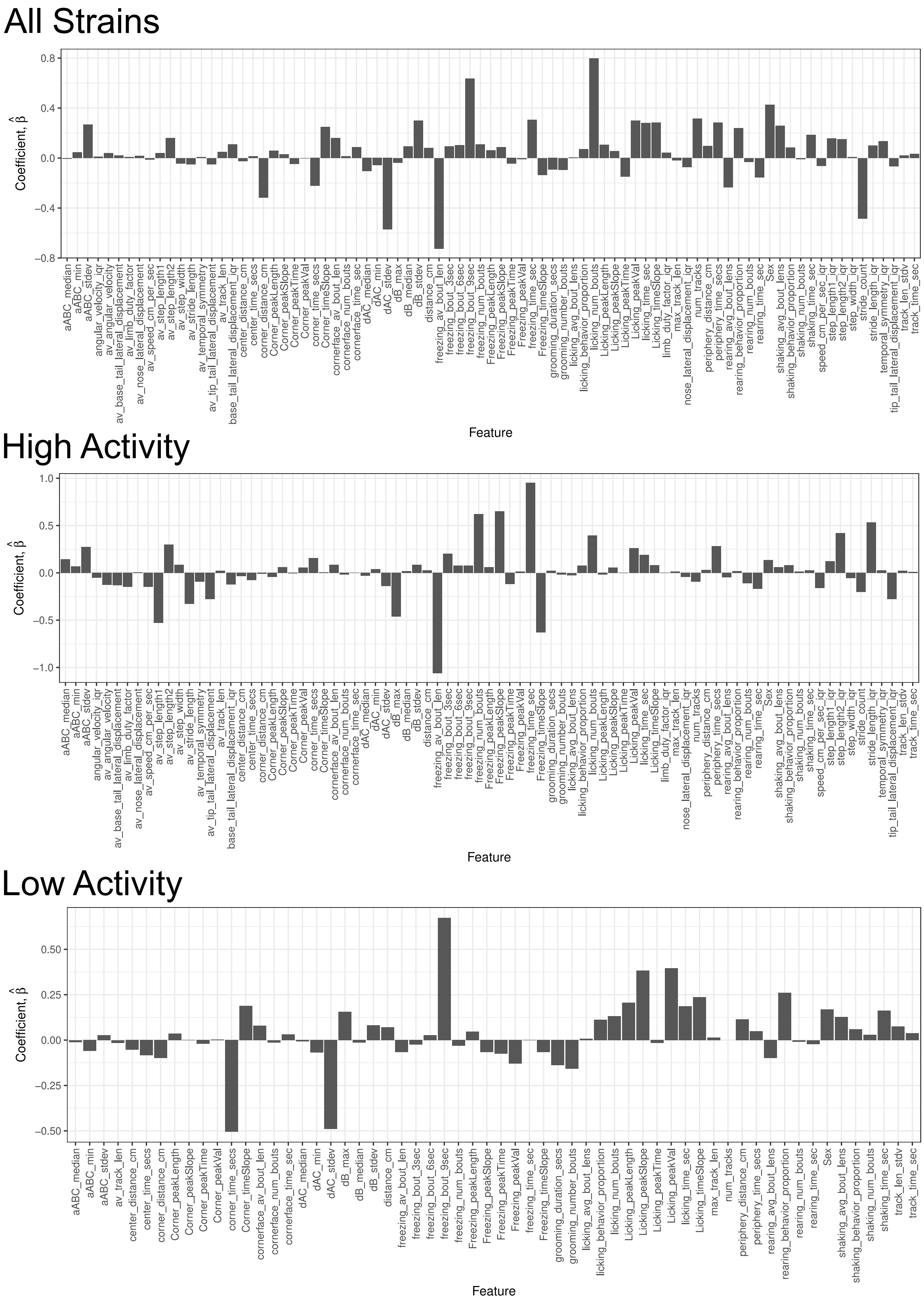


Figure S2: Feature importance for the All strain model, High activity strains model, and low activity strains model is shown. The heights of the bars represent the estimated posterior means, $\hat{\beta}$.

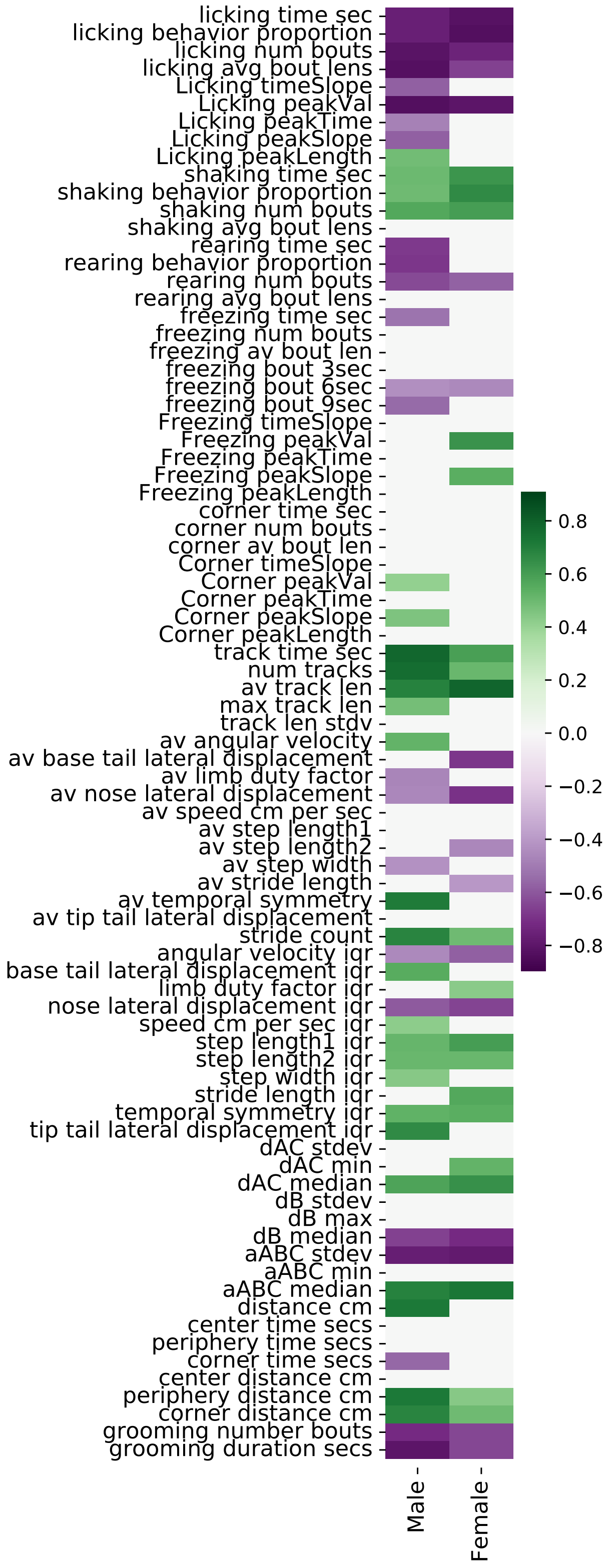


Figure S3: Correlations between feature and morphine dose for male and female C57BL/6J.
